# The landscape of driver mutations in cutaneous squamous cell carcinoma

**DOI:** 10.1101/2020.12.13.422581

**Authors:** Darwin Chang, A. Hunter Shain

## Abstract

Cutaneous squamous cell carcinoma is a form of skin cancer originating from keratinocytes in the skin. It is the second most common type of cancer and is responsible for an estimated 8000 deaths per year in the United States. Compared to other cancer subtypes with similar incidences and death tolls, our understanding of the somatic mutations driving cutaneous squamous cell carcinoma is limited. The main challenge is that these tumors have high mutation burdens, primarily a consequence of UV-radiation-induced DNA damage from sunlight, making it difficult to distinguish driver mutations from passenger mutations. We overcame this challenge by performing a meta-analysis of publicly available sequencing data covering 105 tumors from 10 different studies. Moreover, we eliminated tumors with issues, such as low neoplastic cell content, and from the tumors that passed quality control, we utilized multiple strategies to reveal genes under selection. In total, we nominated 30 cancer genes. Among the more novel genes, mutations frequently affected *EP300*, *PBRM1*, *USP28*, and *CHUK*. Collectively, mutations in the NOTCH and p53 pathways were ubiquitous, and to a lesser extent, mutations affected genes in the Hippo pathway, genes in the Ras/MAPK/PI3K pathway, genes critical for cell-cycle checkpoint control, and genes encoding chromatin remodeling factors. Taken together, our study provides a catalogue of driver genes in cutaneous squamous cell carcinoma, offering points of therapeutic intervention and insights into the biology of cutaneous squamous cell carcinoma.

## Introduction

Over the past decade, large-scale DNA-sequencing studies have profiled a wide range of different cancers^1,2^. These studies have revealed candidate genes for targeted therapy and genetically distinct subtypes of cancer – information that has changed the way in which many cancers are treated. Moreover, at a basic science level, these studies have revealed fundamental insights into the biology of these cancers, often forming the basis of downstream hypothesis-driven work.

Given these achievements, there has been momentum to genomically profile the rarest of cancer subtypes^1,2^, yet cutaneous squamous cell carcinoma, the second most common form of cancer in the United States^3^, has largely been overlooked. Thirty-four cancer subtypes were included in The Cancer Genome Atlas program (TCGA) – a comprehensive effort to catalogue the driver genes in cancer – but regrettably, cutaneous squamous cell carcinoma was left out. Several individual laboratories have sequenced the exomes or genomes of cutaneous squamous cell carcinomas, examples of which are here^4–13^, but the small size of each study and difficulties in interpreting the high mutational loads in cutaneous squamous cell carcinomas have precluded the research community from settling upon a consensus set of driver genes.

One reason why large-scale sequencing consortiums have overlooked cutaneous squamous cell carcinomas is because cutaneous squamous cell carcinomas are thought of as non-life-threatening tumors, however, this reputation is misleading. Most cutaneous squamous cell carcinomas are caught at an early stage, reducing their mortality, but 8000 people per year still die from this disease in the United States^3,14–16^. To put this death toll in perspective, it is on par with that of melanoma^17^, for which nearly 1000 tumors have been sequenced, to date, at exome or genome resolution^18^.

A better understanding of the genetic drivers of cutaneous squamous cell carcinoma promises to improve treatment strategies. The current standard of care is for patients to receive immune-checkpoint blockade therapies, but roughly half do not respond and the responses are not always durable^19^. In addition, the risk of cutaneous squamous cell carcinoma is nearly 100-fold higher in immunosuppressed patients, such as organ transplant recipients, who are typically not eligible to receive immunotherapies^20^. Establishing the driver mutations in cutaneous squamous cell carcinoma promises to reveal new points for therapeutic intervention in this deadly tumor subtype. Towards this goal, we performed a meta-analysis of publicly available exome sequencing data from cutaneous squamous cell carcinomas.

## Results

### Assembling a cohort of cutaneous squamous cell carcinomas

We performed a literature search to identify whole-exome or whole-genome sequencing studies of cutaneous squamous cell carcinoma in which raw sequencing data was made publicly available. In total, we identified 105 tumors spanning 10 studies (Table 1, Table S1)^4–12^. We assessed the quality of sequencing data and removed 17 tumors from subsequent analyses (see methods for exclusion criteria). The remaining 88 tumors were retained, though we accounted for our ability to call somatic mutations in each tumor before comparing them.

**Table 1.** Summary of exome or genome sequencing studies analyzed in this meta-analysis. The number of tumors, listed here, corresponds to unique tumors, whose data was made publicly available, and thus may not match the reported size from the original studies.

| **Studies** | **Sample Size** |
| --- | --- |
| Durinck, S. *et. al.*, *Cancer Disc.*, 2011<sup>9</sup> | 8 |
| Wang, N.J. *et. al.*, *PNAS*, 2011<sup>11</sup> | 4 |
| South, A. P. *et. al.*, *JID*, 2014<sup>10</sup> | 20 |
| Zheng C. L. *et. al.*, *Cell Reports*, 2014<sup>12</sup> | 4 |
| Cammameri *et. al.*, *Nature Comm.*, 2016<sup>8</sup> | 10 |
| Chitsazzadeh *et. al.*, *Nature Comm.*, 2016<sup>6</sup> | 7 |
| Yilmaz, A. S. *et. al.*, *Cancer*, 2017<sup>4</sup> | 6 |
| Cho, R. J. *et. al.*, *Sci. Trans. Med.*, 2018<sup>5</sup> | 27 |
| Inman, G. J. *et. al.*, *Nature Comm*, 2018<sup>7</sup> | 10 |
| Ji, A. L. *et. al.*, *Cell*, 2020<sup>13</sup> | 9 |
| Total | 105 |

The main issues affecting our ability to detect somatic mutations in each tumor were: the neoplastic cell content, the mean sequencing coverage, and/or the variability in sequencing coverage. We bioinformatically quantified tumor cellularities, and they ranged from 12% to 99%. The mean sequencing coverages ranged from 12.4X to 498X. Finally, some tumors had high sequencing coverages, on average, but extreme variability in coverage, primarily linked to the GC content of their targets (see Fig. S1A for an example). To account for each of these potential issues, we used the Footprints software^21^ to count the exact number of basepairs in each sample with sufficient sequencing coverage to make a mutation call. For the average sample, we could detect mutations at 91.2% of target bases, though this ranged from 52.3% to nearly 100% (Fig. S1B).

Establishing the extent to which we could detect mutations in each tumor allowed us to accurately calculate mutation burdens, irrespective of technical variables that are known to distort these measurements^22^. Moreover, the performance of cancer gene discovery tools deteriorates when there are large portions of the exome for which a mutation call cannot be made, and we were able to exclude problematic tumors from these analyses (Fig. S1B). Altogether, we improved the caliber of candidate cancer genes by aggregating a large cohort of tumors, reanalyzing the raw sequencing data, and applying rigorous quality control measures for sample inclusion.

### Subtypes of cutaneous squamous cell carcinoma

We calculated the mutation burden and the proportion of each tumor’s mutations that were attributable to established mutational signatures (Fig. 1), revealing five distinct subtypes of cutaneous squamous cell carcinoma among the tumors analyzed in this study.

**Figure 1.**
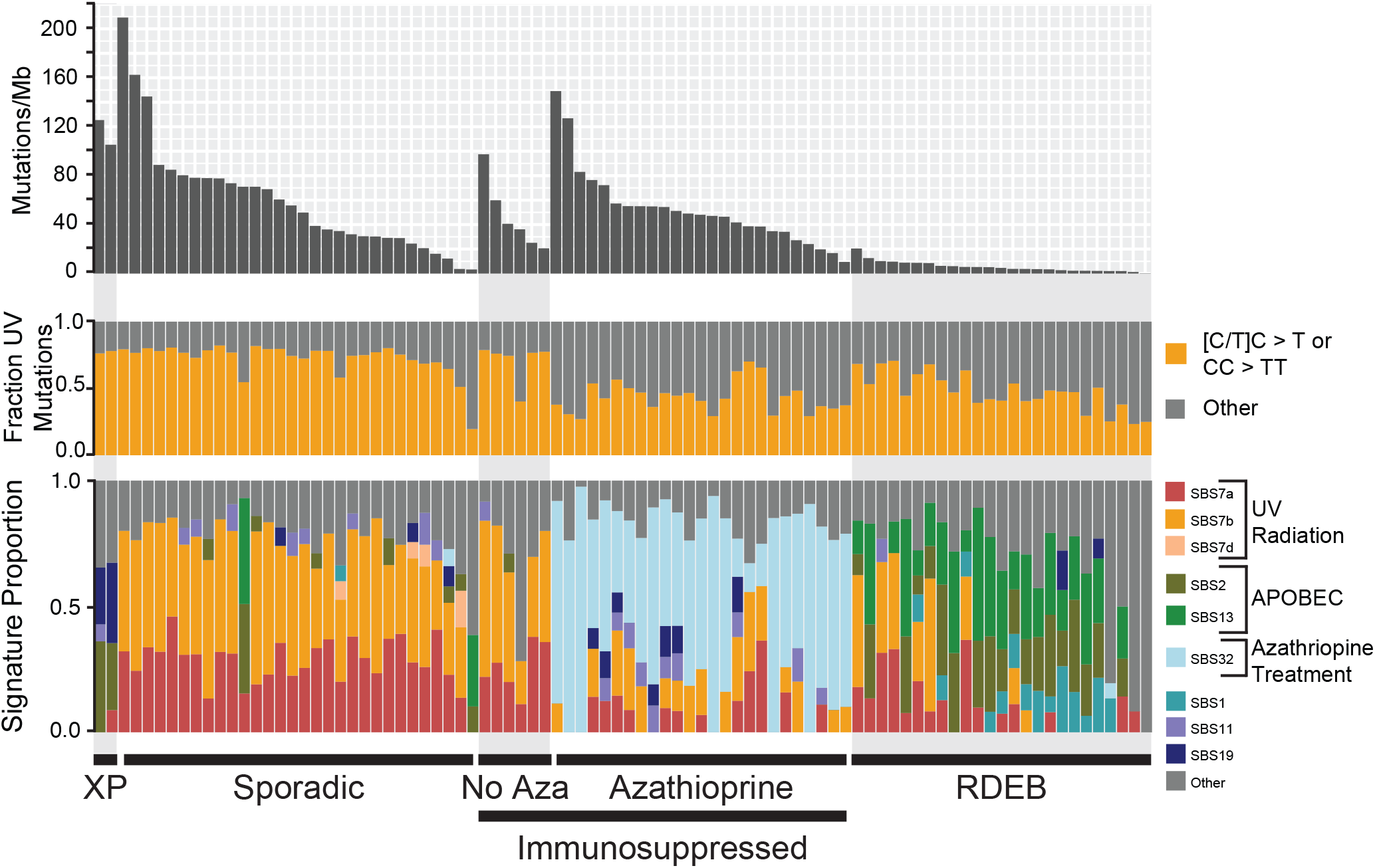
Subtypes of cutaneous squamous cell carcinoma. Each vertical bar corresponds to a single tumor. Top track: The mutation burden (mutations per megabase) of each tumor. Middle track: The fraction of mutations within each tumor matching the canonical UV radiation signature. Bottom track: The fraction of mutations attributable to established mutational signatures within each tumor. The proposed etiology of select signatures is indicated. XP = tumors from patients with xeroderma pigmentosum. Sporadic = tumors from patients with no known comorbidities. Immunosuppressed: tumors from immunosuppressed patients -- this group is further stratified by the usage or absence of azathioprine as an immunosuppressive drug. RDEB = tumors from patients with recessive dystrophic epidermolysis bullosa.

First, two cutaneous squamous cell carcinomas came from patients with xeroderma pigmentosum – a rare hereditary disorder characterized by extreme sensitivity to UV radiation and caused by germline mutations in genes involved in nucleotide excision repair^23^. As expected, these two tumors had high mutation burdens with a high frequency of cytosine to thymine transitions at the 3’ basepairs of consecutive pyrimidines (the classic mutation that arises from UV radiation^24^). Interestingly, they did not have a high proportion of “signature 7” mutations (a mutational signature extracted from pan-cancer analyses and attributed to UV radiation^25,26^). The absence of signature 7 was due to differences in the trinucleotide contexts of mutations arising in wild-type versus mutant-*XPC* tumors (Fig. S2) and illustrates how the repertoire of known mutational signatures remains incomplete.

Second, there were cutaneous squamous cell carcinomas that arose sporadically in patients with no known comorbidities. These cutaneous squamous cell carcinomas had high mutation burdens, and the majority of mutations were attributable to UV radiation (Fig. 1).

Furthermore, there were two distinct types of cutaneous squamous cell carcinomas arising from immunosuppressed patients, who were primarily organ-transplant recipients. As previously reported^7^, cutaneous squamous cell carcinomas from patients treated with Azathioprine (as a means to prevent transplant rejection) had high mutation burdens with high proportions of signature 32 (Fig. 1). Azathioprine increases the risk of cutaneous squamous cell carcinoma beyond the risk conferred by other immunosuppressive agents^27^ because Azathioprine is both immunosuppressive and a potent mutagen^28,29^. Patients on other immunosuppressive drug regimens had comparably lower mutation burdens, primarily attributable to UV radiation (Fig. 1).

Finally, there were cutaneous squamous cell carcinomas from patients with recessive dystrophic epidermolysis bullosa (RDEB), a rare hereditary disorder characterized by chronic blistering in the skin and caused by germline mutations in collagen VII (*COL7A1*). As previously noted^5^, these tumors had relatively low mutation burdens, primarily from APOBEC-mediated mutagenesis (Fig. 1).

### Nomination of driver mutations in cutaneous squamous cell carcinoma

Genes under positive selection in cancer are distinguished by having significantly more mutations than the background mutation rate at that locus would predict^30^. We utilized four cancer gene discovery tools to reveal such genes: MutSig^31^, dN/dS^32^, LOFsigrank^33^, and OncodriveFML^34^. Collectively, these tools nominated 12 genes total, including a subset of 7 genes by at least two tools (Fig. 2, Table S2).

**Figure 2.**
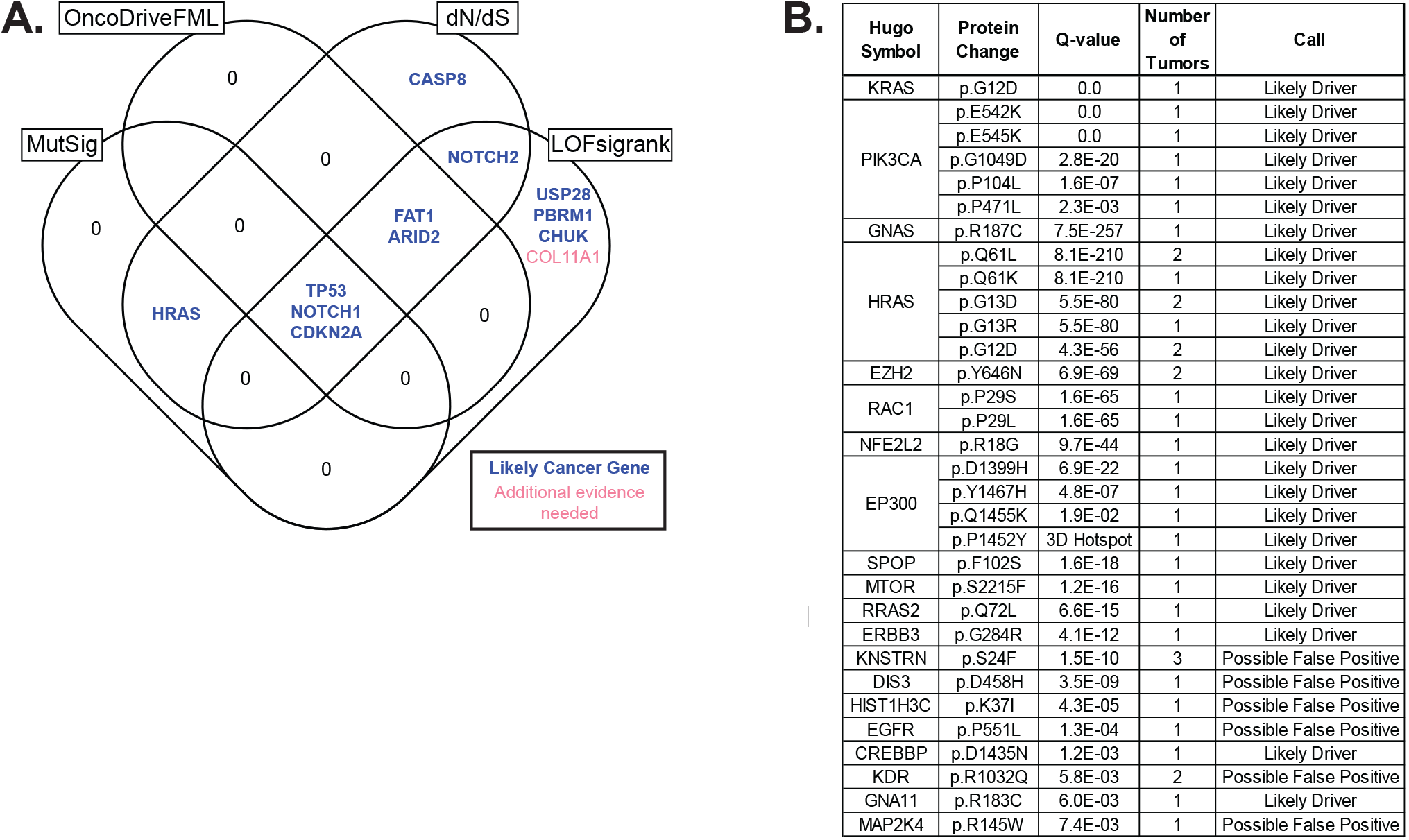
Nomination of cancer genes in cutaneous squamous cell carcinoma. **A.** A Venn diagram depicting nominated cancer genes from four separate cancer gene discovery programs, each designed to identify genes under positive selection in cancer. The set of candidate genes were further curated, as described, to nominate candidates for which additional evidence is warranted (red text) or not (blue text). **B.** A list of mutations in our study that overlap mutations in the cancerhotspot.org database. Mutations are grouped by gene and ordered by their q-values (lowest to highest). These mutations were also curated, as described, to nominate candidates for which additional evidence is warranted (Possible False Positive) or not (Likely Driver).

While these tools nominated a relatively small number of genes, the number of genes nominated is in line with our statistical power to detect cancer genes in this study. It is challenging to identify genes under selection in cutaneous squamous cell carcinoma because of the high background of passenger mutations. Lawrence and colleagues performed a power analysis to determine the detection limit of cancer genes given the mutation burden of the tumor subtype and the number of tumors analyzed^35^. Based on the parameters of our study, we were powered to detect genes under selection if their mutation frequencies were approximately 15% or greater.

To nominate driver mutations that were too infrequent to show evidence of positive selection in this dataset alone, we searched for overlapping mutations in the cancerhotspot.org database (Fig. 2B). This database contains mutations identified from pan-cancer analyses that cluster within genes^36^ – a common pattern for gain-of-function mutations. We reasoned that if a mutation shows evidence of positive selection from pan-cancer analyses, and the exact same mutation was present in our dataset, then it deserves consideration as a driver of cutaneous squamous cell carcinoma.

Finally, we nominated genes with focal copy number alterations. Focal, homozygous deletions affected the *CDKN2A* and *PTEN* tumor suppressor genes (Fig. S3). A focal, heterozygous deletion affected *AJUBA* in one tumor, and in the same tumor there was a point mutation affecting the other allele (Fig. S3B). Focal amplifications affected: *CCND1*, *MDM2*, *YAP1*, and *RAP1B* (Fig. S4). *RAP1B* is a ras-related-protein, and the tumor with amplification of *RAP1B* also had a point mutation affecting the amplified allele. This point mutation was analogous to mutations known to activate other Ras genes (Fig. S4 E,F).

### Removal of false positive driver mutations

The earliest generation of algorithms to discover cancer genes assumed a background mutation rate that is uniform across the genome (an assumption that is not true^37^), resulting in the nomination poor quality candidates in high mutation burden cancers^38,39^. The algorithms, utilized here, have improved background mutation rate models, but the determinants of the mutation rate across the genome are complex and remain incompletely understood^37^, leaving open the possibility of false positives. We curated the nominated genes, as described below, to root out unlikely cancer genes, though we acknowledge that these are ultimately judgement calls.

We concluded that *COL11A1* (Fig. 2A) requires additional evidence to be considered a driver gene. *COL11A1* has a borderline significant q-value (Table S2); it is poorly expressed in the keratinocyte lineage (Fig. S5); and it has no known role in keratinocyte biology. We also determined that *DIS3*, *HIST1H3C*, *KDR*, and *MAP2K4* (Fig. 2B) need additional evidence to be considered driver genes. The hotspot mutations affecting these genes had q-values that were low in comparison to others in the cancerhotspot.org database, and OncoKb, a precision oncology consortium at Memorial Sloan Kettering Cancer Center^40^, classifies these hotspot mutations as unlikely to be oncogenic.

Finally, we do not believe there is sufficient evidence to classify the hotspot mutation affecting *KNSTRN* as oncogenic. The mutation is annotated as coding, but this appears to be based on an erroneous gene model. From RNA-sequencing data of normal skin and cutaneous squamous cell carcinoma, expression of *KNSTRN* begins downstream of the mutation site (Fig. S5B), implying that the mutation affects the promoter of *KNSTRN*. Promoter mutations are ubiquitous in sun-exposed cancers^41,42^ because transcription factors at the promoter can bend DNA in ways that render their binding elements vulnerable to mutagenesis by UV radiation^43^. Regrettably, these types of annotation errors are not uncommon – many hotspot mutations in melanoma, which were initially thought to be coding mutations, were subsequently revealed to be promoter mutations after further studies^18,33,44^. While there is a study suggesting *KNSTRN^S24F^* is oncogenic^45^, the supporting evidence presumes the mutation is coding.

### Novel candidates in cutaneous squamous cell carcinoma

We compared the genes nominated in our meta-analysis to those from eight other studies that have proposed cancer genes in cutaneous squamous cell carcinoma^5–10,46,47^ (Fig. 3). *TP53*, *NOTCH1*, *NOTCH2*, *CDKN2A*, and *HRAS* were proposed by a majority of the other studies and also nominated by us. Moreover, *FAT1*, *ARID2*, *CASP8*, *CREBBP*, *AJUBA*, *PTEN*, *PIK3CA*, *RAC1*, *EZH2*, *KRAS*, *NFE2L2*, and *MTOR* were nominated in 1-3 studies each as well as by us, lending credibility to their pathogenic roles in cutaneous squamous cell carcinoma.

**Figure 3.**
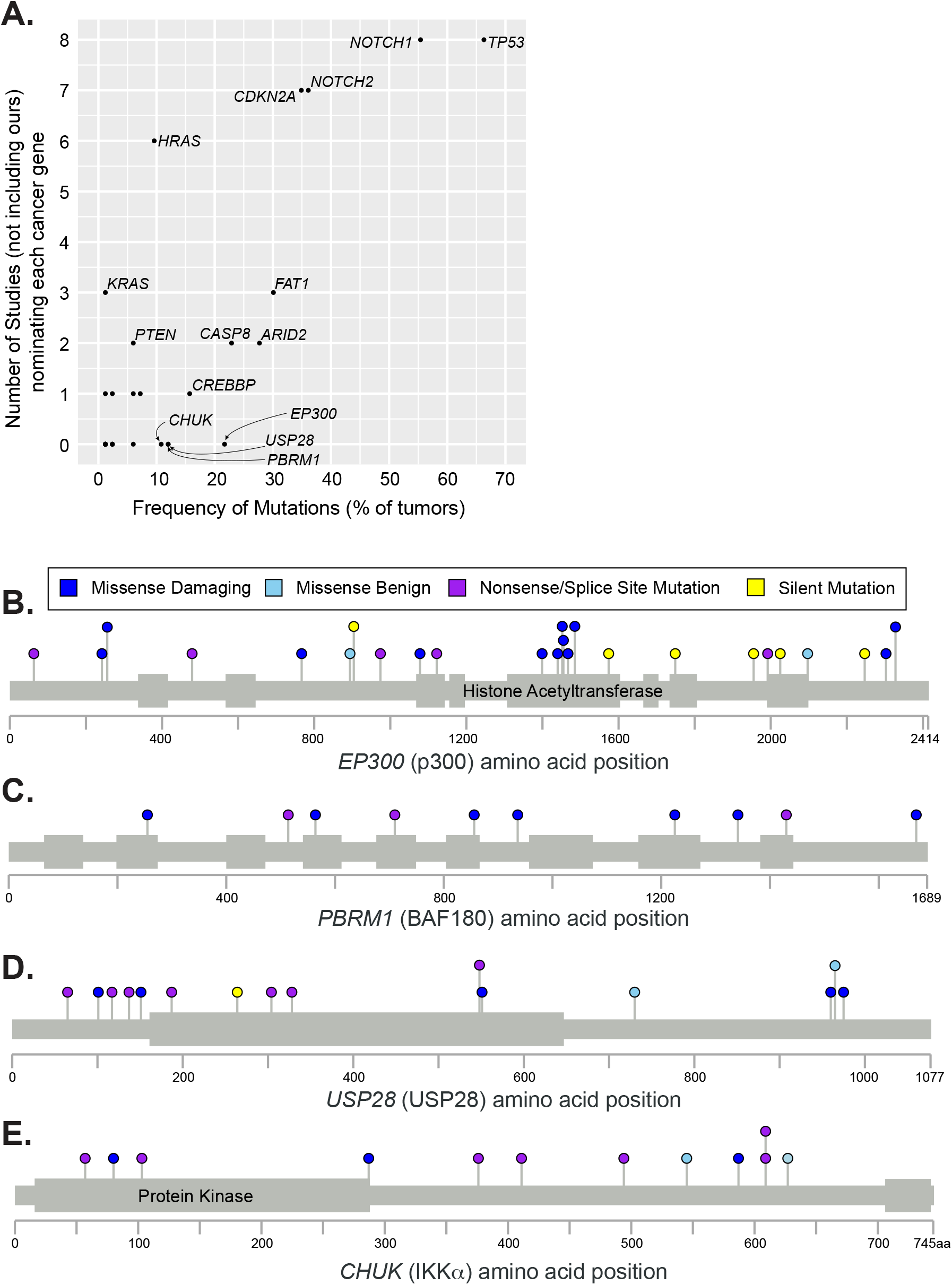
Candidate cancer genes in cutaneous squamous cell carcinoma. **A.** The genes nominated in our meta-analysis are stratified by their mutation frequency (x-axis) and how often they were nominated in 8 previous studies (y-axis) that catalogued drivers of cutaneous squamous cell carcinoma. EP300, PBRM1, USP28, and CHUK were mutated in greater than 10% of tumors but not implicated by other studies. Lollipop diagrams portray the spectrum of mutations in each of these four genes in panels **B-E.**

We nominated 12 genes that were not implicated in the other studies. Many of these genes harbored hotspot mutations that occurred relatively infrequently (see Fig. 2B for the full list), likely explaining why they were not noted in other analyses. However, four genes were mutated in greater than 10% of tumors: *EP300*, *PBRM1*, *USP28*, and *CHUK*.

*EP300* (p300) encodes a histone acetyltransferase that is a critical transcriptional co-activator of NOTCH^48^. *EP300* had frequent loss-of-function mutations, including missense mutations that clustered in the histone acetyltransferase domain (Fig. 3B). Several of these missense mutations have been functionally confirmed to eliminate histone acetyltransferase activity of the protein^49^. *EP300* has also been implicated as a tumor suppressor gene in esophageal squamous cell carcinoma^50^.

*PBRM1* encodes a subunit of the SWI/SNF chromatin remodeling complex and has been implicated as a tumor suppressor gene in a wide range of other cancers^51^. *PBRM1* had deleterious mutations occurring throughout the length of the protein (Fig. 3C). Of note, another member of the SWI/SNF chromatin remodeling complex, *ARID2*, was also implicated as a tumor suppressor gene in cutaneous squamous cell carcinoma.

*USP28* encodes a deubiquitinase that stabilizes key proteins involved in DNA repair^52^. It is required for DNA-damage-induced apoptosis mediated through the Chk2-p53-PUMA pathway^52^. *USP28* was nominated here because of its high frequency of truncating mutations (Fig. 3D).

*CHUK* encodes a protein, also known as IκB Kinase α (IKKα), that is involved in the NFκB signaling pathway. *CHUK* knockout mice are born with thickened skin, and their cutaneous keratinocytes are unable to differentiate, resulting in death shortly after birth^53^. An identical phenotype has been observed in humans with a defective *CHUK* gene^54^. Collectively, these studies implicate *CHUK* as a key factor governing growth and differentiation of keratinocytes in skin. In cutaneous squamous cell carcinoma, *CHUK* had a high frequency of truncating mutations, and somatic alterations affected both alleles in most tumors (Fig. 3E).

We also checked for genes nominated in other studies but not by our analyses. *KMT2D* was the only gene implicated in more than one of the other studies interrogated here (Fig. S6A). The majority of *KMT2D* mutations were silent or missense mutations predicted to be benign (Fig. S6B). Even under relaxed thresholds of significance (Table S2), *KMT2D* would not have been nominated here, though future studies may reveal more compelling evidence of selection.

### Recurrent pathways disrupted in cutaneous squamous cell carcinoma

The individual genes, nominated here, encode proteins that participate in a core set of signaling pathways perturbed in cutaneous squamous cell carcinoma (Fig. 4). Mutations in genes encoding proteins involved in the NOTCH and p53 pathways were ubiquitous in cutaneous squamous cell carcinomas. The NOTCH pathway had loss-of-function mutations occurring in 80% of tumors, and the p53 pathway had loss-of-function mutations occurring in 71% of tumors. Mutations in these pathways appear to be defining features of cutaneous squamous cell carcinoma.

**Figure. 4.**
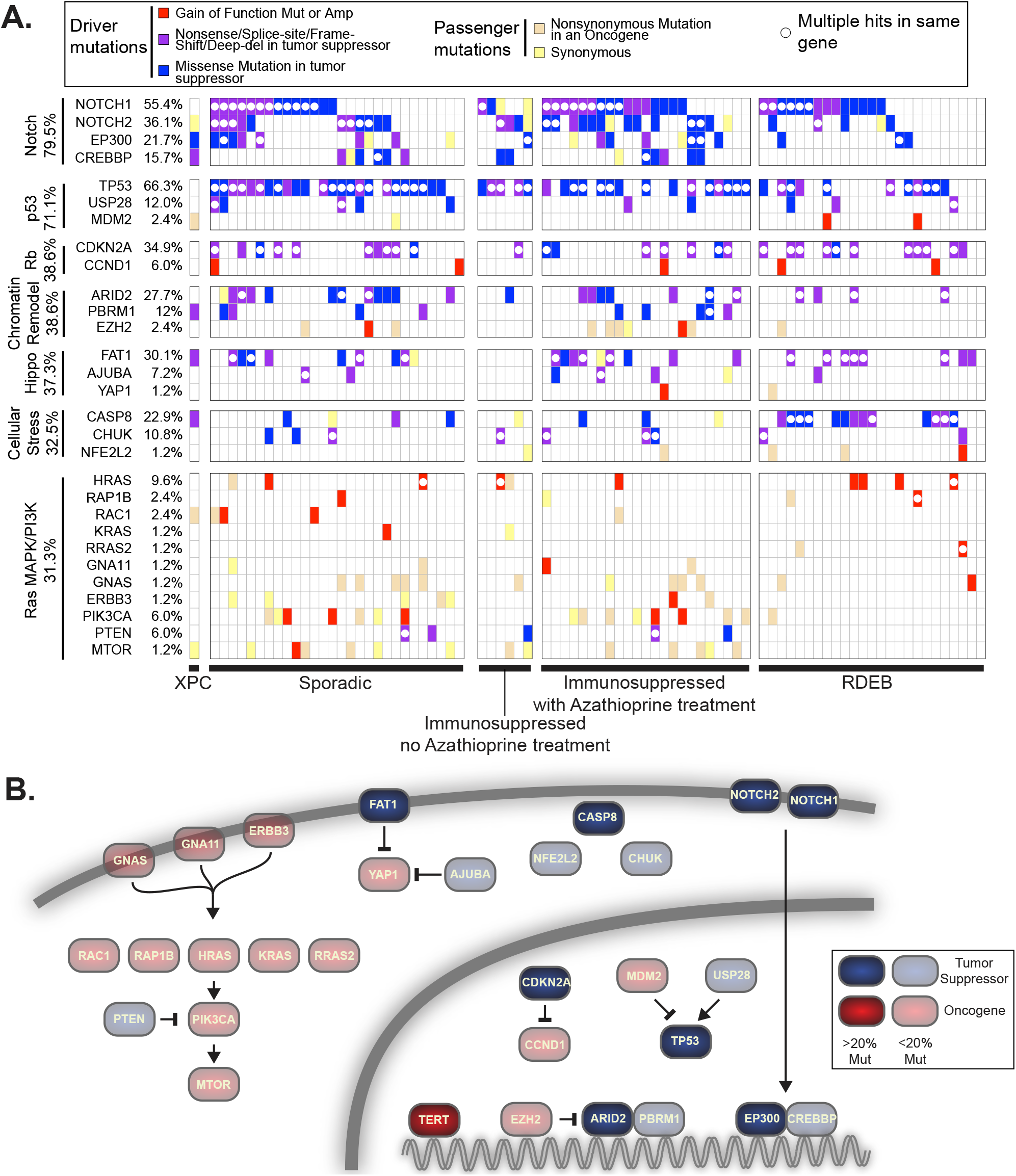
The landscape of driver mutations in cutaneous squamous cell carcinoma. **A.** Tiling plot of the genetic alterations (rows) in each tumor (columns). Genes and tumors are further organized into pathways and clinical subtypes. The percentages of samples harboring pathogenic alterations are indicated. Mut, mutation; Amp, amplification; XPC, xeroderma pigmentosum; RDEB, recessive dystrophic epidermolysis bullosa. **B.** Pathways affected. TERT was not implicated in this study, which focused on coding mutations, but is included here because it is known to have promoter mutations, as described.

Other pathways were recurrently disrupted, albeit to a lesser extent. Mutations that disrupt cell-cycle-checkpoint control occurred in 39% of tumors, primarily affecting the *CDKN2A* gene. Mutations that disrupt the SWI/SNF chromatin remodeling complex occurred in 38% of tumors. Mutations that activate the Hippo pathway occurred in 37% of tumors. We broadly grouped together *CASP8*, *CHUK*, and *NFE2L2*, which were collectively mutated in 33% of tumors. These genes mediate cellular responses to stress, such as inflammation and oxidative stress. More work will be needed to determine whether and how these genes are related. Finally, mutations that activate the Mitogen Activated Protein Kinase (MAPK) and/or PhosphoInositide 3-Kinase (PI3K) pathways occurred in 31% of tumors.

We next interrogated whether mutations affecting specific genes, pathways, or tumor subtypes overlapped more or less than would be expected by chance. Cutaneous squamous cell carcinomas from patients with recessive dystrophic epidermolysis bullosa (RDEB) were enriched with mutations affecting *CASP8* (Fig. 4A, Table S3). CASP8 mediates cellular apoptosis in response to inflammatory cytokines^55,56^. Skin from patients with RDEB is chronically blistering and inflamed, likely explaining the selective pressure to accumulate *CASP8* mutations in this subtype of cutaneous squamous cell carcinoma. Other comparisons did not reach statistical significance after accounting for multiple hypothesis testing (see Table S3 for a complete list of comparisons).

Most of the studies in this meta-analysis were exome-sequencing studies, prohibiting us from analyzing mutations in non-coding portions of the genome. *TERT* promoter mutations are common in many cancer subtypes, and while we were unable to investigate the locus, other studies have reported a high frequency of *TERT* promoter mutations in cutaneous squamous cell carcinoma^57^, prompting us to include *TERT* among our final list of cancer genes (Fig. 4B). Future studies will be needed to more systematically interrogate the role of non-coding mutations in cutaneous squamous cell carcinoma.

## Discussion

In this meta-analysis of exome sequencing data, we analyzed the largest cohort of cutaneous squamous cell carcinomas to date, upheld rigorous quality control standards for sample inclusion, and utilized state-of-the-art cancer gene discovery algorithms to nominate cancer genes. In total, we nominated 30 cancer genes, known to operate in a core set of signaling pathways, that were perturbed in cutaneous squamous cell carcinoma. Our study suggests new cancer genes and helps clarify which candidates from previous studies are likely *bona fide* driver genes in cutaneous squamous cell carcinoma.

Future work is still needed to understand the driver genes in cutaneous squamous cell carcinoma. Cancer gene discovery studies have likely reached a saturation point for many cancers, but this is not the case for cutaneous squamous cell carcinoma. Despite the size of our meta-analysis, we could only detect cancer genes with mutations in 15% of more of tumors. We overcame this limitation, in part, by identifying genes with well-characterized hotspot mutations and/or focal copy number alterations; however, there are likely many cancer genes in cutaneous squamous cell carcinoma that have yet to be discovered. Our study also focused on the exome, prohibiting us from identifying driver mutations in non-coding regions of the genome or from identifying structural variants and viral integrations that may play a pathogenic role.

Taken together, our study provides the most detailed catalogue of driver genes in cutaneous squamous cell carcinoma to date, offers points of therapeutic vulnerability, and reveals critical insights into the basic biology of cutaneous squamous cell carcinoma.

## Methods

### Selection of studies

We performed a literature search to identify whole-exome or whole-genome sequencing studies of cutaneous squamous cell carcinoma that made their raw sequencing data publicly available as of September 1^st^, 2020. The studies meeting this inclusion criteria are summarized in Table 1. The number of samples shown in Table 1 may not match the reported numbers in each study because some studies re-analyzed previously published data, or we were unable to retrieve the entirety of the raw sequencing dataset.

### Removal of 17 samples

We assessed the quality of sequencing data and removed 17 tumors from all analyses. Thirteen of these tumors had few, if any, discernible point mutations, and among the point mutations detected, their mutant allele frequencies were close to our detection limit. These patterns suggest poor sampling of the neoplastic cells. Two cases had less than five-fold coverage in the reference tissues, making it difficult to confidently distinguish somatic mutations from germline single nucleotide polymorphisms (SNPs). One tumor and reference pair were not properly matched, which was evident from their patterns of germline SNPs. Finally, one reference tissue had high levels of tumor contamination, prohibiting us from sensitively detecting somatic mutations.

### Calling somatic point mutations

We collected either fastq or bam files from each study. Fastq files underwent quality checks using FastQC and were subsequently aligned to the hg19 reference genome using the BWA-MEM algorithm (v0.7.13). These were further groomed and deduplicated using Genome Analysis Toolkit (v4.1.2.0) and Picard (v4.1.2.0).

Somatic point mutations were called using Mutect2 (v4.1.2.0) by comparing each tumor bam to a corresponding reference bam, thus producing an initial set of candidate somatic mutations. The variants were annotated using Funcotator (v4.1.2.0) and further filtered to remove suspected sequencing artifacts, extremely subclonal mutations, and/or mutations from unrelated clones of keratinocytes. In parallel, indels were called using Pindel (v0.2.5) and further filtered. Our filtering scripts are available here: https://github.com/darwinchangz/ShainMutectFilter.

To provide an overview, the script uses samtools mpileup to count the number of reference and mutant reads for each variant. Variants with low overall coverage were removed, and variants with few supporting mutant reads were also removed. Finally, we calculated tumor cellularity in each sample, and removed variants that were not predicted to be in at least 40% of tumor cells. The main reason we removed these variants is because it was difficult to distinguish whether they were from subclones within the tumor or from unrelated clones of mutant keratinocytes. Normal skin is comprised of clones of keratinocytes, many of which harbor pathogenic mutations^6,58^. We have observed that these clones commingle with adjacent skin tumors and are often unintentionally included in microdissections^59^.

### Calling Heterozygous SNPs

We also identified a high-confidence set of germline heterozygous SNPs from the reference bams corresponding to each patient. Knowing these SNPs allowed us to measure allelic imbalance, thereby revealing tumor cellularity (detailed below) and corroborating copy number alterations within tumors. To identify heterozygous, germline SNPs, we called variants in the reference tissue as compared to the reference genome using FreeBayes (v1.3.1). Next, we filtered these variants to include only those that overlapped known 1000 genomes sites and which had 40-60% variant allele frequency (VAF).

### Inferring Tumor Cellularity

We used multiple methods, if possible, to infer the neoplastic cell content from each tumor. The methods used for each tumor are listed in Table S1 and further described below.

“Allelic Imbalance of SNPs over Deletions”: We calculated tumor cellularity from the degree of allelic imbalance of heterozygous, germline SNPs over chromosomal arms with deletions in the tumor. This strategy assumes the deletions are fully clonal and there remains only one copy of the remaining chromosome in each tumor cell. A deletion results in a complete loss of an allele within the tumor cells. As a result, sequencing reads from the deleted allele are assumed to come from non-neoplastic cells. Tumor cellularity can therefore be calculated from ratio of reads mapping to the A and B alleles as described^59^.

“Allelic Imbalance of SNPs over Copy Number Neutral LOH”: Similar to the above strategy, we calculated tumor cellularity from the degree of allelic imbalance of heterozygous, germline SNPs over chromosomal arms with copy-number-neutral loss-of-heterozygosity (LOH). This strategy assumes that copy number neutral LOH is fully clonal and there are two copies of the remaining allele in each tumor cell. Copy number neutral LOH results in complete loss of an allele within the tumor cells. As a result, sequencing reads mapping to the lost allele are assumed to come from the non-neoplastic cells. Tumor cellularity can therefore be calculated from ratio of reads mapping to the A and B alleles as described^59^.

“Modal Somatic MAF”: In addition to investigating the variant allele frequencies of heterozygous, germline SNPs, we also used the mutant allele frequencies (MAFs) of somatic mutations. The mutant allele frequency of a somatic mutation that is fully-clonal and heterozygous should be 50%, but will decrease with stromal contamination. For each tumor, we plotted a histogram of mutant allele frequencies and determined the “peak” or “modal” mutant allele frequency, and we doubled these values to infer tumor cellularity.

“Median Somatic MAF”: For a small number of tumors, the density of somatic mutations was insufficient to produce a smooth histogram. In these cases, we determined the median MAF from all somatic mutations, and we doubled this value to infer tumor cellularity. If the patient was male, we separately calculated the median MAF of somatic mutations on the sex chromosomes and incorporated these values without doubling.

### Determining statistical power to call somatic mutations (related to figure S1)

To identify a somatic mutation, there must be sufficient coverage in both the reference and the tumor. Therefore, for each tumor/reference pair, we calculated the footprint for which sequencing coverage was sufficient to call somatic mutations.

In the reference, sufficient coverage is necessary to detect both alleles, thus ensuring that a variant in the tumor is a somatic mutation and not a germline SNP. We required at least 6-fold coverage in the reference to call a somatic mutation. Assuming each allele is randomly sampled during sequencing, the probability of both alleles being sampled at least once with 6-fold coverage is 96.9% (two-tailed binomial test). We used the Footprints software^21^ to calculate the precise number of base-pairs that achieved 6-fold coverage (or greater) in each reference bam and designated this value as the “call-able” footprint for each reference bam (Table S1).

In the tumor, there needs to be sufficient coverage to detect the mutant allele. We required our somatic mutation calls to have at least 4 unique reads. Some mutation callers, including MuTect2, which was used in this study, will attempt to call mutations with fewer reads, but in practice, we found those calls to be of poor quality and filtered them out. We considered a site to be “call-able” if it had 8-fold effective tumor coverage. “Effective tumor coverage” refers to the sequencing coverage derived from the tumor after discounting the proportion of reads from non-tumor cells. For example, if a tumor sample has 100-fold total coverage and 8% tumor cellularity, then the effective tumor coverage would be 8-fold. Assuming that the alleles are randomly sampled during sequencing, their relative coverages will fit a binomial distribution and 8-fold effective tumor coverage is sufficient to call a heterozygous somatic mutation 50% of the time (two-tailed binomial test). We used the Footprints software^21^ to calculate the precise number of base-pairs that achieved 8-fold effective tumor coverage or greater in each tumor bam and designated this value as the “call-able” footprint for each tumor bam (Table S1).

For each tumor/reference pair, we took the minimum “call-able” footprint between the tumor and the reference and designated that value to be the “call-able” footprint for that sample. We subsequently divided the “call-able” footprint by the bait territory that was targeted to indicate the fraction of target basepairs for which we were statistically powered to recognize mutations. These numbers are reported in figure S1A.

There were primarily three variables that reduced statistical power to recognize mutations: 1. Low overall coverage, 2. Low tumor cellularity, 3. Extreme variability in coverage (e.g. from GC-selection biases introduced during hybridization).

### Calling Copy Number Alterations

Copy number alterations were inferred from the DNA-sequencing data using CNVkit^60^. CNVkit can be run in reference or reference-free mode. We elected to run CNVkit in reference mode using the panel of normals from each study. This approach consistently produced the least noisy copy number profiles, as compared to reference-free mode or a universal reference. All other parameters were run on their default settings.

### Calculating tumor mutation burden and inferring mutational signature (related to figure 1)

When calling somatic point mutations, we only considered mutations that were estimated to be in at least 40% of tumor cells. This was helpful in comparing the mutation burdens from tumors across the different studies for which there was considerable variability in sequencing coverages. High-sequencing coverage permits the detection of subclonal mutations, which would artificially inflate the mutation burden of a tumor, compared to another with lower coverage, if subclonal mutations are counted. There were also differences in our ability to detect clonal mutations in each tumor (described in more detail in the “*Determining statistical power to call somatic mutations*” section above). To address this issue, we divided the number of clonal mutations in each tumor by the footprint which we were statistically powered to detect mutations in each tumor.

To perform mutational signature analysis, surrounding genomic contexts were applied to single nucleotide variants identified in each clone using the Biostrings hg19 human genome sequence package (BSgenome.Hsapiens.UCSC.hg19 v1.4.0). Variant contexts were used to assess the proportion of each clone’s mutational landscape that could be attributed to a mutagenic process using the deconstructSigs R package (v1.9.0). A set of 48 signatures recently described^26^ were analysed. The results of these analyses are shown in the “Signature Proportion” stacked barplot of figure 1. In parallel, we performed a simpler analysis of dinucleotide contexts to identify cytosine to thymine transitions at the 3’ basepair of dipyrimidines or cytosine-cytosine to thymine-thymine mutations (see the “UV” column of Table S4) – these are the classic mutation types attributed to UV radiation, and the results are shown in the “Fraction UV Mutations” stacked barplot of figure 1.

### Nomination of driver genes

We used four cancer gene discovery programs to nominate cancer genes: MutSig^31^, LOFsigrank^33^, dN/dS^32^, and OncodriveFML^34^. dN/dS was run in covariate value mode, which combines the synonymous mutations in a gene with its epigenomic covariates to determine the background mutation rate. The other programs were run on their default settings. All genes with q-values of less than 0.2 are shown in Table S2. For the purposes of this study, a gene was considered significant if its q-value was less than 0.05 (Figure 2).

Next, we cross-referenced the cancerhotspot.org database to identify mutations found in our study. We included mutations with a q-value of less than 0.01, but relaxed this threshold for all secondary hotspots within a gene. For example, EP300 had two hotspot mutations with q-values of 6.9E-22 and 4.8E-07, but we also show additional hotspots in the vicinity, one with a q-value of 1.9E-02 and one that was a predicted hotspot using the 3D hotspot algorithm.

Finally, driver genes were curated as described in the main text to root out potential false positives.

### RNA-Sequencing analysis (related to figure S5)

Two of the studies analyzed in this meta-analysis had RNA-sequencing data available^5,6^. These datasets covered both normal skin (n=17 samples) and cutaneous squamous cell carcinoma (n=17 samples). We downloaded the raw sequencing data, aligned with STAR, and quantified gene expression with RSEM, as previously described^61^. In figure S5, we show the Fragment Per Kilobase of transcript per Million reads (FPKM) values for candidate genes (see figure 2 for a list of candidates). FPKM values normalize for gene length and read depth, allowing the comparison of gene expression levels across genes. In figure S5, we combined RNA-sequencing reads from all 34 samples over the *KNSTRN* gene, demonstrating that the mutant hostpot is not expressed.

### Mutational overlap analysis (related to Table S3)

We interrogated whether mutations affecting specific genes, pathways, or tumor subtypes overlapped more or less than would be expected by chance. We restricted our analyses to genes, pathways, or tumor subtypes with at least 16 mutant tumors – the minimum number that could reach statistical significance with our sample size. P-values for individual comparisons were calculated using the Fisher’s exact test. We corrected for multiple hypothesis testing by computing false discovery rates (q-values) using the Benjamini-Hochberg procedure. A full list of p- and q-values for each comparison is shown in Table S3.

## Figure and Table legends

**Table S1. A summary of samples included in this study and their sequencing metrics.** We identified 105 cutaneous squamous cell carcinomas from 101 patients (4 patients had multiple tumors) for which sequencing data was publicly available. This table summarizes available information pertaining to these patients and their tumors as well as relevant sequencing metrics. Note that some tumors had whole genome sequencing, but we restricted our analyses to the exome. Methods to infer tumor cellularity are explained in more detail under *Inferring Tumor Cellularity* in Methods. RDEB = Recessive Dystrophic Epidermolysis Bullosa. XPC = Xeroderma Pigmentosum.

**Table S2. Genes nominated to be under positive selection in cutaneous squamous cell carcinoma.** All genes nominated by each of the four cancer gene discovery tools with a q-value below 0.2 are shown. For the purposes of this study, we considered a gene to significant if it had a q-value of less than 0.05. In the first tab, we show genes nominated from an analysis of all tumors passing quality control, and in the subsequent tabs, we show the genes nominated from analysis of clinically distinct types of cutaneous squamous cell carcinoma.

**Table S3. Genes, pathways, and tumor subtypes with significantly overlapping (or non-overlapping) mutations.** P-values reflect the results of a two-tailed Fisher exact test, and q-values account for multiple hypothesis testing using the Benjamini-Hochberg procedure.

**Table S4. Somatic mutations in cutaneous squamous cell carcinoma.** Somatic mutations that passed filtering were annotated using Funcotator and shown here.

## Acknowledgements

We wish to acknowledgement funding support from the University of California Cancer Research Coordinating Committee, the UCSF Resource Allocation Program (Mt. Zion Health Fund), the UCSF Department of Dermatology, and the National Cancer Institute (K22 award).

---

**Figure S1.**
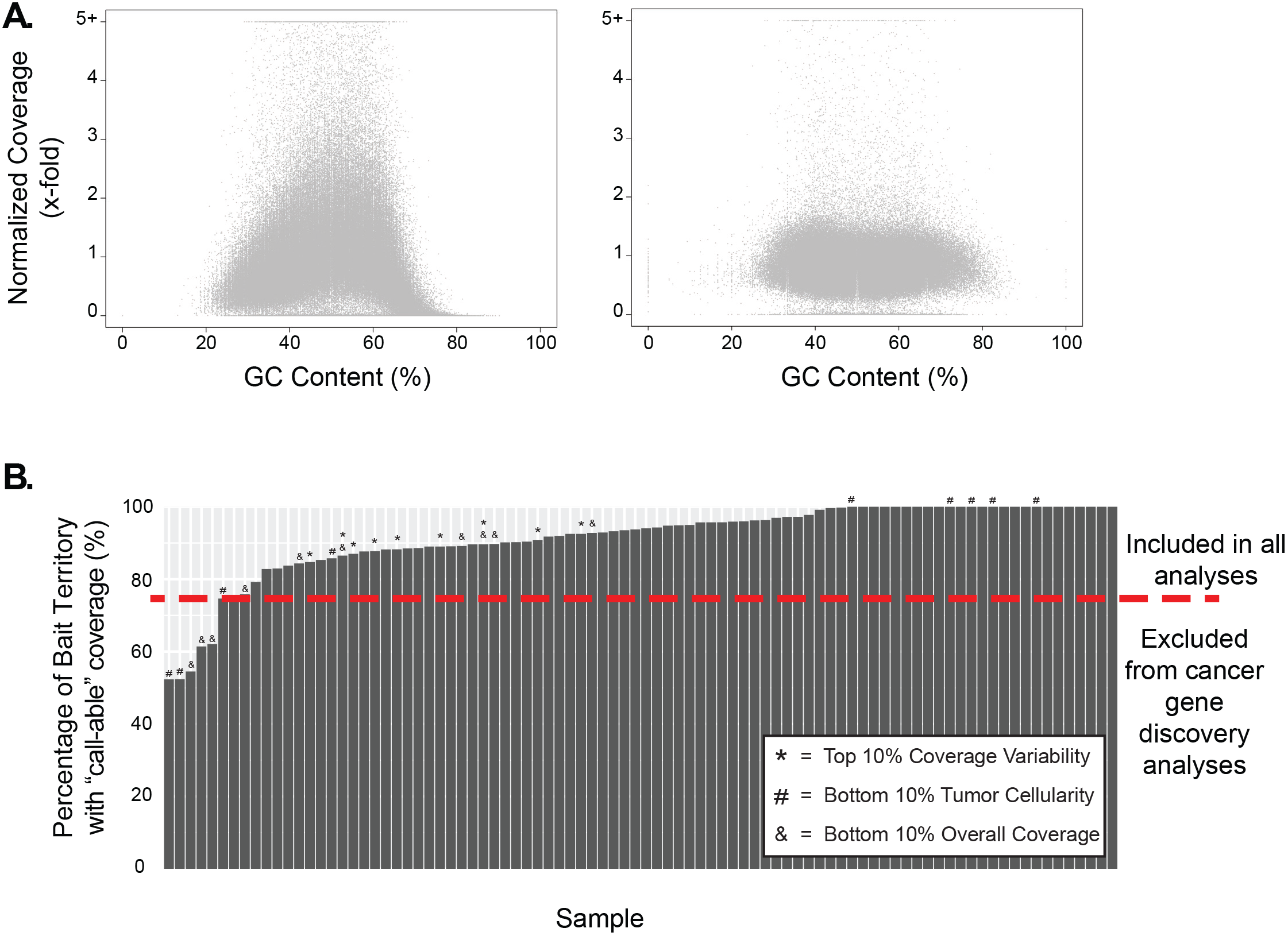
Ability to detect mutations in each tumor varies substantially. **A.** Some tumors had high overall coverage but extreme variability in coverage, primarily linked to GC content, thus diminishing our ability to detect mutations within large portions of the bait territory. An example of a tumor in the top 10% of coverage variability (left panel) and a tumor in the bottom 10% of coverage variability (right panel). Each datapoint corresponds to a bait interval, stratified by its GC content and sample-normalized coverage. **B**. The percentage of base pairs within the target bait territory with sufficient coverage to call a mutation. Tumors with low overall sequencing coverage, extreme variability in sequencing coverage, and/or low tumor cellularity tended to have a lower proportion of “call-able” mutations. Tumors with greater than 75% “call-able” mutations were included in all analyses, whereas those with fewer were only included in certain analyses, as described.

**Figure S2.**
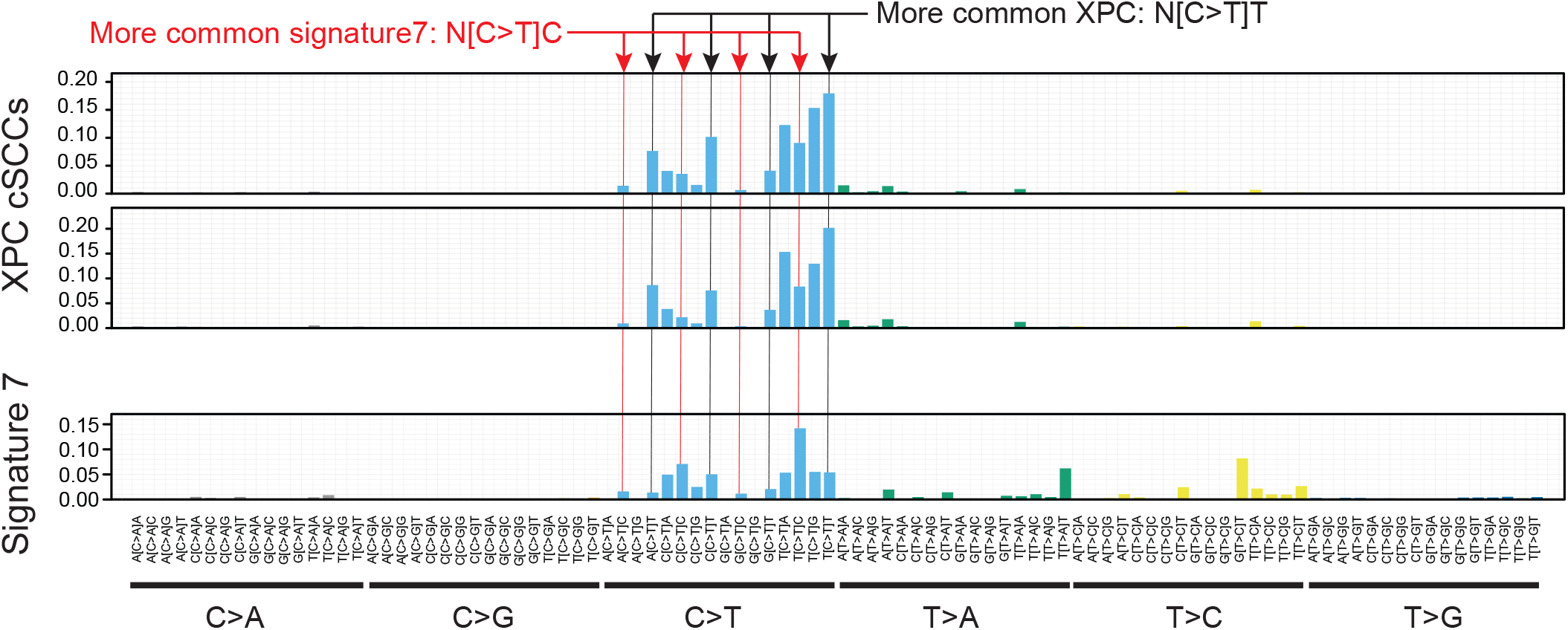
Somatic mutations in Xeroderma Pigmentosum tumors occur in a distinct tri-nucleotide context. The tri-nucleotide context of mutations in two XPC−/− tumors as compared to signature 7. C>T transitions at dipyrimidines were common in both, however the basepair downstream (three prime) to the mutation site differed with thymines more common in XPC−/− tumors and cytosines more common in signature 7.

**Figure S3.**
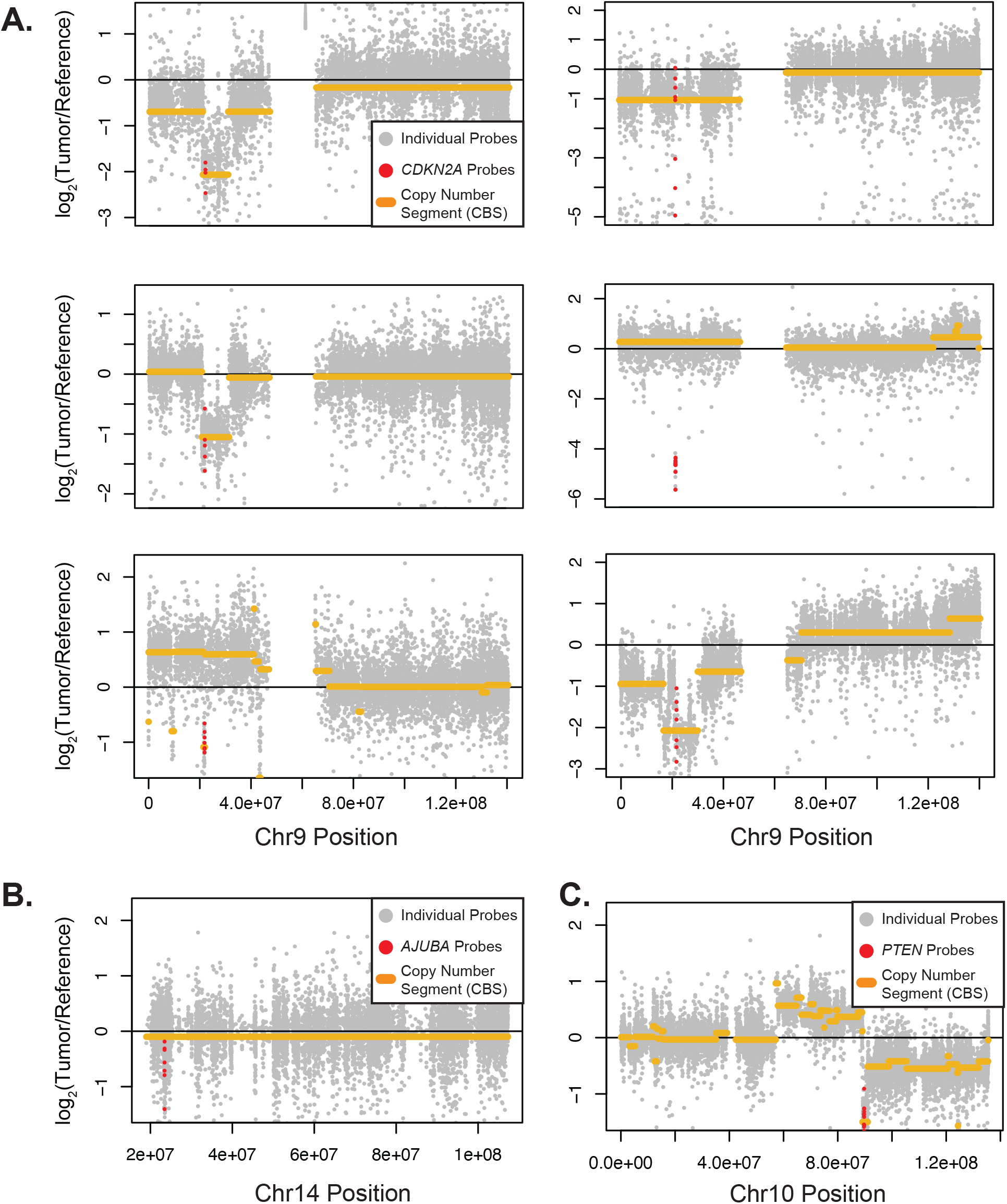
Focal deletions affecting CDKN2A, AJUBA, and PTEN. Copy number over individual genomic positions (data points) with copy number segments (yellow lines) overlaid. Colored data points correspond to probes covering CDKN2A (panel A), AJUBA (panel B), and PTEN (panel C).

**Figure S4.**
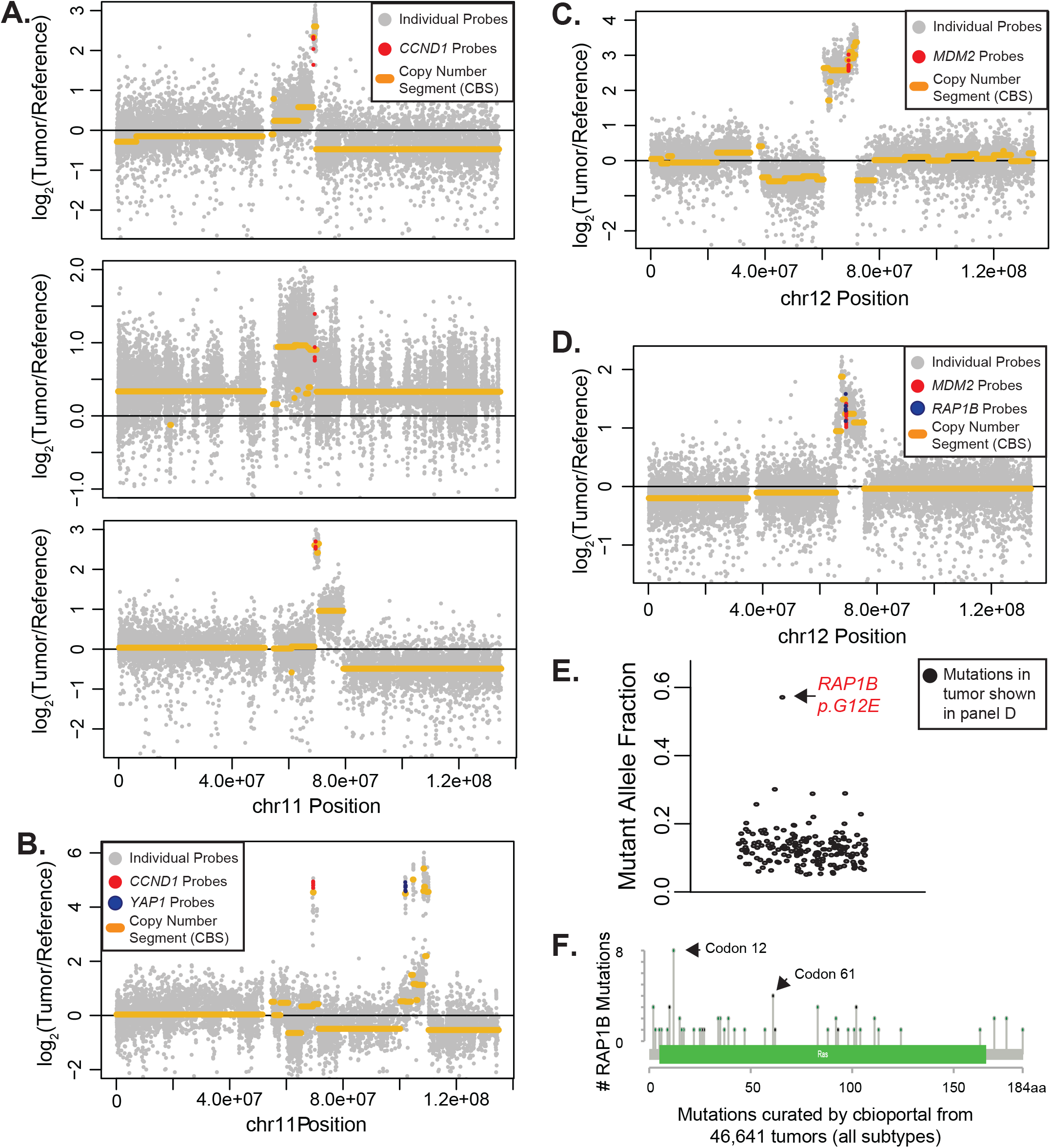
Focal amplification affecting CCCND1, YAP1, MDM2, and/or RAP1B in six tumors. Copy number over individual genomic positions (data points) with copy number segments (yellow lines) overlaid. Colored data points correspond to probes covering CCND1 (panels A and B), YAP1 (panel B), MDM2 (panels C and D), and RAP1B (panel D). Note that amplifications converge upon the noted genes across these six tumors. The tumor with amplification of RAP1B (shown in panel D) also had a point mutation affecting the 12th codon of RAP1B (panel E). The mutant allele frequency of this point mutation was higher than all other mutations in that tumor, consistent with the mutation affecting the amplified allele. RAP1B is a ras-like protein with recurrent mutations affecting codons 12 and 61 across cancer (panel F), analogous to mutations known to activate other ras-like proteins.

**Figure S5.**
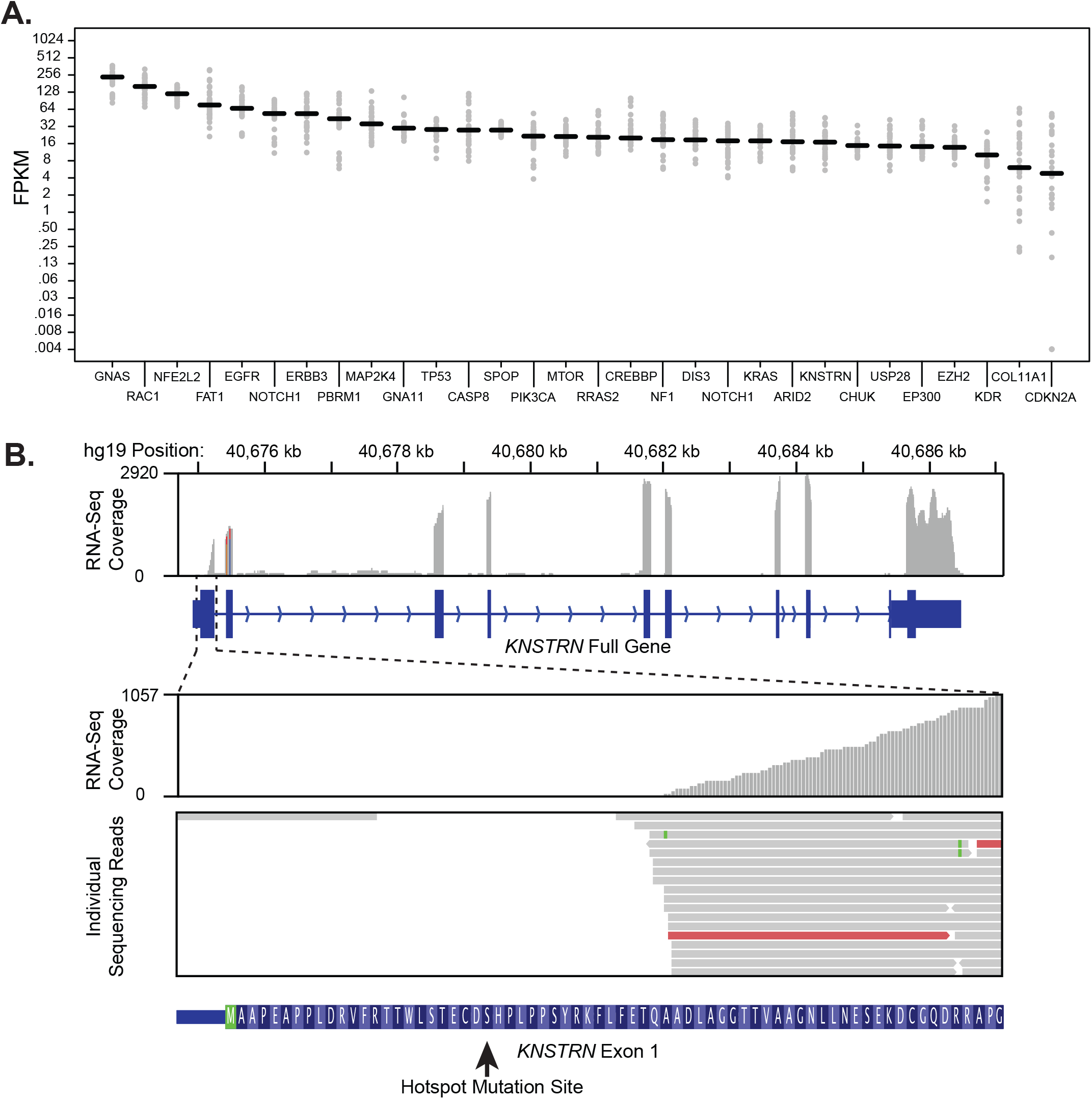
Gene expression of candidate genes. **A.** The Fragments Per Kilobase of transcript per Million reads (FPKM) values -- a measurement that normalizes for both gene length and sequencing depth in RNA-sequencing data -- of candidate genes. RNA-sequencing data derived from both normal skin samples (n=17 samples) and cutaneous squamous cell carcinomas (n=17 tumors). This data was used to flag potential false positive candidates based on their poor expression in normal and neoplastic keratinocytes. **B.** Transcription of KNSTRN begins downstream of the hotspot mutation site. RNA-sequencing data was aggregated from normal skin (n=17) and cutaneous squamous cell carcinoma (n=17). Top panel -- RNA-sequencing coverage over the KNSTRN gene. Coverage was approximately 2,000 to 3,000-fold over most exons with the exception of exon1. Bottom panel -- A zoomed inset of RNA-sequencing coverage over exon1. While sequencing coverage surpassed 1,000-fold at the three-prime junction of exon1, there were no reads spanning the hotspot mutation site from the any of the 34 samples, in-line with our prediction that the mutation is non-coding.

**Figure S6.**
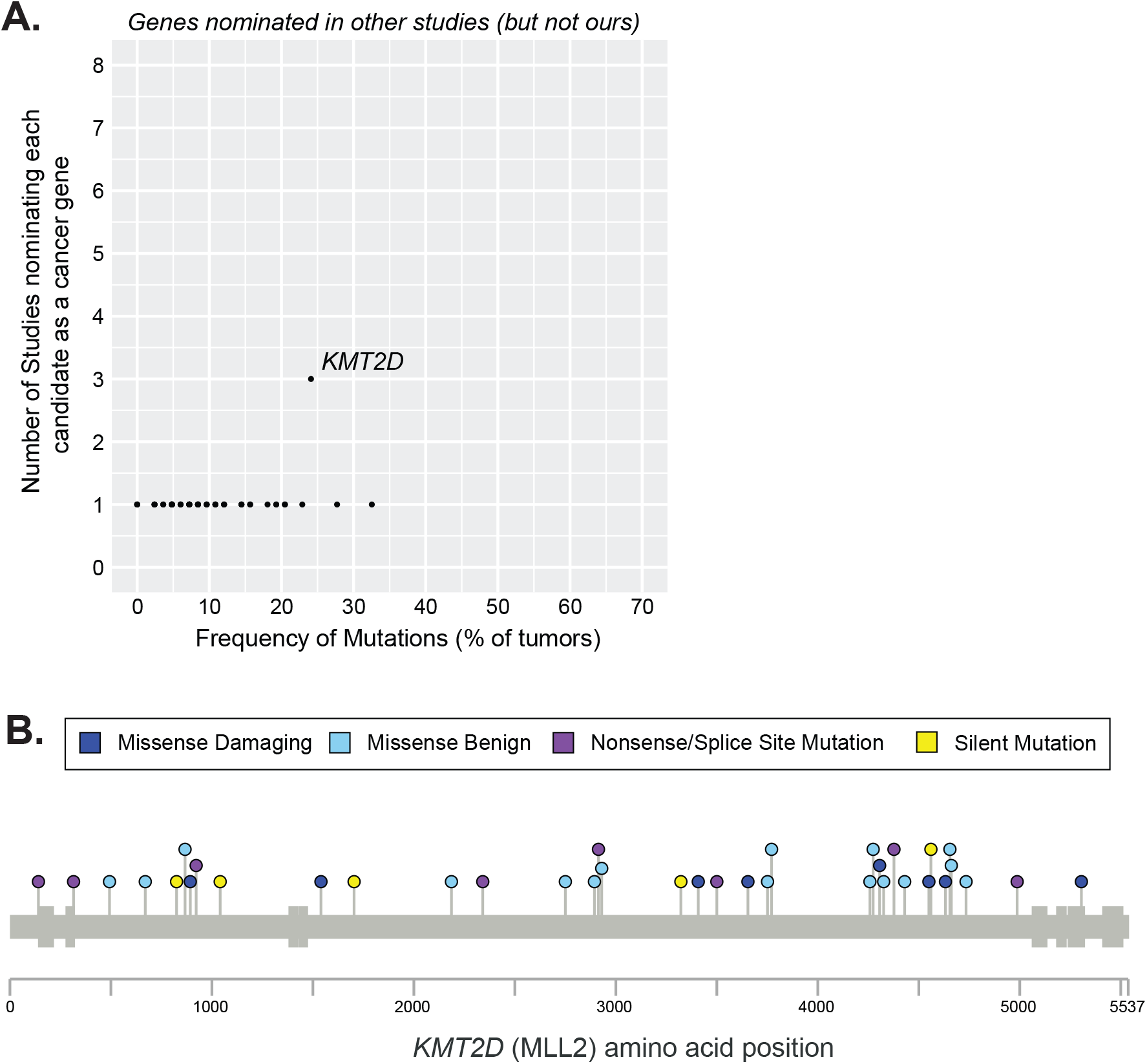
KMT2D has a benign spectrum of mutations. **A.** Genes nominated by other studies, but not our study, are stratified by their mutation frequency (x-axis) and how often they were nominated in 8 previous studies (y-axis) that catalogued drivers of cutaneous squamous cell carcinoma. KMT2D was the only gene recurrently implicated in other studies but not ours. **B.** Lollipop diagram portrays the spectrum of mutations. KMT2D was not nominated by cancer gene discovery algorithms because of its high frequency of silent mutations and missense mutations predicted to be benign.

## References

1. Garraway, L. A. & Lander, E. S. Lessons from the cancer genome. Cell 153, 17–37 (2013).

2. Vogelstein, B. et al. Cancer genome landscapes. Science 339, 1546–1558 (2013).

3. Nehal, K. S. & Bichakjian, C. K. Update on Keratinocyte Carcinomas. N. Engl. J. Med. 379, 363–374 (2018).

4. Yilmaz, A. S. et al. Differential mutation frequencies in metastatic cutaneous squamous cell carcinomas versus primary tumors. Cancer 123, 1184–1193 (2017).

5. Cho, R. J. et al. APOBEC mutation drives early-onset squamous cell carcinomas in recessive dystrophic epidermolysis bullosa. Sci. Transl. Med. 10, (2018).

6. Chitsazzadeh, V. et al. Cross-species identification of genomic drivers of squamous cell carcinoma development across preneoplastic intermediates. Nat. Commun. 7, 12601 (2016).

7. Inman, G. J. et al. The genomic landscape of cutaneous SCC reveals drivers and a novel azathioprine associated mutational signature. Nat. Commun. 9, 3667 (2018).

8. Cammareri, P. et al. Inactivation of TGFβ receptors in stem cells drives cutaneous squamous cell carcinoma. Nat. Commun. 7, 12493 (2016).

9. Durinck, S. et al. Temporal dissection of tumorigenesis in primary cancers. Cancer Discov. 1, 137–143 (2011).

10. South, A. P. et al. NOTCH1 mutations occur early during cutaneous squamous cell carcinogenesis. J. Invest. Dermatol. 134, 2630–2638 (2014).

11. Wang, N. J. et al. Loss-of-function mutations in Notch receptors in cutaneous and lung squamous cell carcinoma. Proc. Natl. Acad. Sci. U. S. A. 108, 17761–17766 (2011).

12. Zheng, C. L. et al. Transcription restores DNA repair to heterochromatin, determining regional mutation rates in cancer genomes. Cell Rep. 9, 1228–1234 (2014).

13. Ji, A. L. et al. Multimodal Analysis of Composition and Spatial Architecture in Human Squamous Cell Carcinoma. Cell 182, 1661–1662 (2020).

14. Karia, P. S., Han, J. & Schmults, C. D. Cutaneous squamous cell carcinoma: estimated incidence of disease, nodal metastasis, and deaths from disease in the United States, 2012. J. Am. Acad. Dermatol. 68, 957–966 (2013).

15. Wu, W. & Weinstock, M. A. Trends of keratinocyte carcinoma mortality rates in the United States as reported on death certificates, 1999 through 2010. Dermatol. Surg. Off. Publ. Am. Soc. Dermatol. Surg. Al 40, 1395–1401 (2014).

16. Mansouri, B. & Housewright, C. D. The Treatment of Actinic Keratoses-The Rule Rather Than the Exception. JAMA Dermatol. 153, 1200 (2017).

17. Siegel, R. L., Miller, K. D. & Jemal, A. Cancer statistics, 2016. CA. Cancer J. Clin. 66, 7–30 (2016).

18. Multi-omic analysis reveals significantly mutated genes and DDX3X as a sex-specific tumor suppressor in cutaneous melanoma | Nature Cancer. https://www.nature.com/articles/s43018-020-0077-8.

19. Migden, M. R. et al. PD-1 Blockade with Cemiplimab in Advanced Cutaneous Squamous-Cell Carcinoma. N. Engl. J. Med. 379, 341–351 (2018).

20. Madeleine, M. M. et al. Epidemiology of keratinocyte carcinomas after organ transplantation. Br. J. Dermatol. 177, 1208–1216 (2017).

21. Tang, J. et al. The genomic landscapes of individual melanocytes from human skin. bioRxiv 2020.03.01.971820 (2020) doi:10.1101/2020.03.01.971820.

22. Stenzinger, A. et al. Tumor mutational burden standardization initiatives: Recommendations for consistent tumor mutational burden assessment in clinical samples to guide immunotherapy treatment decisions. Genes. Chromosomes Cancer 58, 578–588 (2019).

23. DiGiovanna, J. J. & Kraemer, K. H. Shining a light on xeroderma pigmentosum. J. Invest. Dermatol. 132, 785–796 (2012).

24. Brash, D. E. UV Signature Mutations. Photochem. Photobiol. 91, 15–26 (2015).

25. Alexandrov, L. B. et al. Signatures of mutational processes in human cancer. Nature 500, 415–421 (2013).

26. Alexandrov, L. B. et al. The repertoire of mutational signatures in human cancer. Nature 578, 94–101 (2020).

27. Jiyad, Z., Olsen, C. M., Burke, M. T., Isbel, N. M. & Green, A. C. Azathioprine and Risk of Skin Cancer in Organ Transplant Recipients: Systematic Review and Meta-Analysis. Am. J. Transplant. Off. J. Am. Soc. Transplant. Am. Soc. Transpl. Surg. 16, 3490–3503 (2016).

28. Harwood, C. A. et al. PTCH mutations in basal cell carcinomas from azathioprine-treated organ transplant recipients. Br. J. Cancer 99, 1276–1284 (2008).

29. O’Donovan, P. et al. Azathioprine and UVA light generate mutagenic oxidative DNA damage. Science 309, 1871–1874 (2005).

30. Martínez-Jiménez, F. et al. A compendium of mutational cancer driver genes. Nat. Rev. Cancer (2020) doi:10.1038/s41568-020-0290-x.

31. Lawrence, M. S. et al. Mutational heterogeneity in cancer and the search for new cancer-associated genes. Nature 499, 214–218 (2013).

32. Martincorena, I. et al. Universal Patterns of Selection in Cancer and Somatic Tissues. Cell 171, 1029–1041.e21 (2017).

33. Shain, A. H. et al. Exome sequencing of desmoplastic melanoma identifies recurrent NFKBIE promoter mutations and diverse activating mutations in the MAPK pathway. Nat. Genet. 47, 1194–1199 (2015).

34. Mularoni, L., Sabarinathan, R., Deu-Pons, J., Gonzalez-Perez, A. & López-Bigas, N. OncodriveFML: a general framework to identify coding and non-coding regions with cancer driver mutations. Genome Biol. 17, 128 (2016).

35. Lawrence, M. S. et al. Discovery and saturation analysis of cancer genes across 21 tumour types. Nature 505, 495–501 (2014).

36. Chang, M. T. et al. Accelerating Discovery of Functional Mutant Alleles in Cancer. Cancer Discov. 8, 174–183 (2018).

37. Gonzalez-Perez, A., Sabarinathan, R. & Lopez-Bigas, N. Local Determinants of the Mutational Landscape of the Human Genome. Cell 177, 101–114 (2019).

38. Shain, A. H. & Bastian, B. C. Raising the bar for melanoma cancer gene discovery. Pigment Cell Melanoma Res. 25, 708–709 (2012).

39. Hodis, E. et al. A Landscape of Driver Mutations in Melanoma. Cell 150, 251–263 (2012).

40. Chakravarty, D. et al. OncoKB: A Precision Oncology Knowledge Base. JCO Precis. Oncol. 2017, (2017).

41. Sabarinathan, R., Mularoni, L., Deu-Pons, J., Gonzalez-Perez, A. & López-Bigas, N. Nucleotide excision repair is impaired by binding of transcription factors to DNA. Nature 532, 264–267 (2016).

42. Perera, D. et al. Differential DNA repair underlies mutation hotspots at active promoters in cancer genomes. Nature 532, 259–263 (2016).

43. Mao, P. et al. ETS transcription factors induce a unique UV damage signature that drives recurrent mutagenesis in melanoma. Nat. Commun. 9, 2626 (2018).

44. Rodríguez-Martínez, M. et al. Evidence That STK19 Is Not an NRAS-dependent Melanoma Driver. Cell 181, 1395–1405.e11 (2020).

45. Lee, C. S. et al. Recurrent point mutations in the kinetochore gene KNSTRN in cutaneous squamous cell carcinoma. Nat. Genet. 46, 1060–1062 (2014).

46. Pickering, C. R. et al. Mutational landscape of aggressive cutaneous squamous cell carcinoma. Clin. Cancer Res. Off. J. Am. Assoc. Cancer Res. 20, 6582–6592 (2014).

47. Li, Y. Y. et al. Genomic analysis of metastatic cutaneous squamous cell carcinoma. Clin. Cancer Res. Off. J. Am. Assoc. Cancer Res. 21, 1447–1456 (2015).

48. Oswald, F. et al. p300 acts as a transcriptional coactivator for mammalian Notch-1. Mol. Cell. Biol. 21, 7761–7774 (2001).

49. Duex, J. E. et al. Functional Impact of Chromatin Remodeling Gene Mutations and Predictive Signature for Therapeutic Response in Bladder Cancer. Mol. Cancer Res. MCR 16, 69–77 (2018).

50. Gao, Y.-B. et al. Genetic landscape of esophageal squamous cell carcinoma. Nat. Genet. 46, 1097–1102 (2014).

51. Shain, A. H. & Pollack, J. R. The spectrum of SWI/SNF mutations, ubiquitous in human cancers. PloS One 8, e55119 (2013).

52. Zhang, D., Zaugg, K., Mak, T. W. & Elledge, S. J. A role for the deubiquitinating enzyme USP28 in control of the DNA-damage response. Cell 126, 529–542 (2006).

53. Li, Q. et al. IKK1-deficient mice exhibit abnormal development of skin and skeleton. Genes Dev. 13, 1322–1328 (1999).

54. Lahtela, J. et al. Mutant CHUK and severe fetal encasement malformation. N. Engl. J. Med. 363, 1631–1637 (2010).

55. Boldin, M. P., Goncharov, T. M., Goltsev, Y. V. & Wallach, D. Involvement of MACH, a novel MORT1/FADD-interacting protease, in Fas/APO-1- and TNF receptor-induced cell death. Cell 85, 803–815 (1996).

56. Muzio, M. et al. FLICE, a novel FADD-homologous ICE/CED-3-like protease, is recruited to the CD95 (Fas/APO-1) death--inducing signaling complex. Cell 85, 817–827 (1996).

57. Griewank, K. G. et al. TERT promoter mutations are frequent in cutaneous basal cell carcinoma and squamous cell carcinoma. PloS One 8, e80354 (2013).

58. Martincorena, I. et al. Tumor evolution. High burden and pervasive positive selection of somatic mutations in normal human skin. Science 348, 880–886 (2015).

59. Shain, A. H. et al. The Genetic Evolution of Melanoma from Precursor Lesions. N. Engl. J. Med. 373, 1926–1936 (2015).

60. Talevich, E., Shain, A. H., Botton, T. & Bastian, B. C. CNVkit: Genome-Wide Copy Number Detection and Visualization from Targeted DNA Sequencing. PLoS Comput. Biol. 12, e1004873 (2016).

61. Shain, A. H. et al. Genomic and Transcriptomic Analysis Reveals Incremental Disruption of Key Signaling Pathways during Melanoma Evolution. Cancer Cell 34, 45–55.e4 (2018).

